# Spatial distribution of hand-grasp motor task activity in spinal cord functional magnetic resonance imaging

**DOI:** 10.1101/2023.04.25.537883

**Authors:** Kimberly J. Hemmerling, Mark A. Hoggarth, Milap S. Sandhu, Todd B. Parrish, Molly G. Bright

**Author notes:** **Corresponding Author:** Kimberly J. Hemmerling 645 N. Michigan Ave., Suite 1100 Chicago, IL 60611 USA.

## Abstract

Upper extremity motor paradigms during spinal cord functional magnetic resonance imaging (fMRI) can provide insight into the functional organization of the cord. Hand-grasping is an important daily function with clinical significance, but previous studies of similar squeezing movements have not reported consistent areas of activity and are limited by sample size and simplistic analysis methods. Here, we study spinal cord fMRI activation using a unimanual isometric hand-grasping task that is calibrated to participant maximum voluntary contraction (MVC). Two task modeling methods were considered: (1) a task regressor derived from an idealized block design (Ideal) and (2) a task regressor based on the recorded force trace normalized to individual MVC (%MVC). Across these two methods, group motor activity was highly lateralized to the hemicord ipsilateral to the side of the task. Activation spanned C5-C8 and was primarily localized to the C7 spinal cord segment. Specific differences in spatial distribution are also observed, such as an increase in C8 and dorsal cord activity when using the %MVC regressor. Furthermore, we explored the impact of data quantity and spatial smoothing on sensitivity to hand-grasp motor task activation. This analysis shows a large increase in number of active voxels associated with the number of fMRI runs, sample size, and spatial smoothing, demonstrating the impact of experimental design choices on motor activation.

## 1. Introduction

Spinal cord functional magnetic resonance imaging (fMRI) offers unique potential to noninvasively visualize functional activation patterns related to sensorimotor activity in the human spinal cord. This not only provides insight into healthy human functional anatomy, but also has the potential to improve our understanding of pathological spinal cord function in injury or disease. fMRI of the human spinal cord was performed as early as 1996^1^, not long after the introduction of brain fMRI in 1990^2^. However, the growth of spinal cord fMRI is disproportionately small relative to its critical importance in the central nervous system. In recent review articles, only about 100-115 articles on spinal cord fMRI were identified, depending on literature search parameters^3, 4^

This slow growth in spinal cord fMRI is likely attributable to its technical challenges. The small cross-sectional size of the cord directly imposes a tradeoff between larger voxels with higher signal-to-noise ratio (SNR) and smaller voxels with higher spatial resolution and reduced partial volume effects. The spinal cord is also surrounded by structures, such as the vertebrae, that vary in magnetic susceptibility and lead to inhomogeneities in the magnetic field. Nearby respiratory structures vary in magnetic susceptibility during breathing, leading to spatiotemporal variation in the main magnetic field^5^. The nearby cardiac-linked pulsatile flow of cerebrospinal fluid (CSF) surrounding the cord additionally confounds fMRI signals^6, 7^. However, several pivotal developments in acquisition and analysis techniques addressing these issues have enabled successes in the spinal cord fMRI field (see Kinany et al. 2022 for a review of these strategies^8^), making this important technique more feasible for translational research.

Since the inception of spinal cord fMRI, motor tasks have been implemented to study the functional organization of the cord^1^, and such tasks have been consistently used since then^9–30^. Upper extremity motor paradigms are the most common; these tasks are feasible in the scanner environment without excessive movement of the trunk and neck. Additionally, the cervical and upper thoracic spinal cord segments, where associated motor activity would be expected, can be imaged with a head/neck coil used in typical brain imaging setups^8^. Motor tasks have been used to characterize aspects of movements in the wrist^12, 24, 26–29^, fingers^12, 16–18, 20–23, 26, 27^, hand^1, 9–11, 15, 19, 30^, ankle/foot^14^, and tongue^13^. It is relevant to note that, although these studies employ motor tasks, some are specifically focused on development or comparison of spinal cord fMRI acquisition techniques^15, 20, 25, 28^, rather than characterizing the activation itself.

In these motor task studies, the spatial distribution of activation in the spinal cord is commonly characterized by its laterality, ventrodorsal distribution, and/or activation peak. Unilateral tasks have frequently demonstrated lateralization of motor activity to the hemicord ipsilateral to the task side^1, 9, 12, 17, 19, 21, 23, 24, 27, 29, 30^, but in other studies laterality was unclear or not present^10, 14, 16, 19, 20^. Weber et al. (2016), stressing the lack of studies providing quantitative evidence of lateralization, demonstrates reliable lateralization to the ipsilateral left and right hemicords with an isometric wrist flexion task^24^. Although motor activation is typically expected in the ventral horn of the spinal cord gray matter, where motor neuron cell bodies are located^31^, activation is not usually confined to this area in spinal cord fMRI studies. Frequently, dorsal horn activity is attributed to some unavoidable effect of proprioceptive and/or sensory feedback coupled with the task^10, 14, 24, 26, 30^. Additionally, activity in the intermediate zone (the gray matter between ventral/dorsal horns) has been ascribed to combined neural processing of unilateral tasks^24^.

Based on anatomical myotomes, each motor task is expected to elicit fMRI activity in specific spinal cord levels. The localization of activity along the rostrocaudal axis is typically described as a peak of motor activity in 1-2 segments^3^. Studies specifically probing rostrocaudal organization utilize several upper limb tasks to elicit distinguishable clusters of motor activity in spinal cord segments^12, 26^. In an early implementation of spinal cord fMRI, Madi et al. (2001) showed alignment of tasks to the predicted anatomical myotome^12^. Building on these findings, Kinany et al. (2019) demonstrated distinct rostrocaudal organization between three upper extremity motor tasks in a larger cohort and linked these findings to electromyography-based spinal mapping^26^.

Here, we aim to evaluate the spatial distribution of spinal cord activation associated with hand grasping, which is of particular importance in numerous pathologies influencing quality of life. For example, abnormal muscle coactivation patterns are associated with impaired grasping ability following stroke^32^, and grip strength may even predict future health outcomes in healthy older adults^33^. In order for spinal cord fMRI to be a valuable tool in understanding such neural pathologies, it is important to establish a normative reference of hand-grasping activity in healthy adults.

Prior spinal cord fMRI studies have used various grasping movements such as squeezing a ball^9, 11, 19, 30^ or fist clenching^10, 15, 25^ **(Supplementary Table S1)**. Ipsilateral activation was not consistently reported and there was no consensus localization of activation along the rostrocaudal axis. Overall, few participants were included in these studies (median N=11), and all but one was published prior to 2010. Image acquisition and processing protocols considered common now (e.g., ZOOMit selective excitation, Spinal Cord Toolbox functions for spatial normalization), were not yet widely accessible or adopted. Early grasping studies establishing the field of spinal cord fMRI focused on subject-level analyses of signal change. After a >10-year gap, modern preprocessing, subject-, and group-level analyses were used in a study using a fist-clenching task, however, the focus of this study was on MR acquisition methods rather than mapping task activation^25^. This exemplifies a long standing concern in fMRI, whereby studies (and particularly early studies) have few participants and report more qualitative subject-level results without robust group statistics^3^, or perform underpowered group analyses that inflate activation estimates and bias our interpretation of activation patterns^34^. Therefore, the neural activation during hand-grasping is not yet well characterized in the human spinal cord and we are equipped to bring modern advances in acquisition and analysis to better understand this question.

In this study, we investigate the spinal cord blood oxygenation level-dependent (BOLD) fMRI response to an isometric left and right unimanual hand-grasping task using state-of-the-art acquisition and analysis methods in a larger cohort of healthy adults (N=30). To minimize variability in the hand grasping task, we target a grasping force relative to an individual’s maximum voluntary contraction (MVC). Targeting is achieved via real-time visual feedback of grasp force achieved by the participant, which is also recorded, allowing flexibility in modeling of the BOLD response to the intended task (i.e., idealized block design or actual %MVC time course). We expect to observe peak motor activity in the ventral horn of the C7-C8^31^ spinal cord segments, ipsilateral to the task side. However, motor activation is not expected to be confined only to this region. Our primary goal is to create a robust and detailed map of hand-grasping spinal cord activation in healthy adults. We characterize the spatial distribution of activation throughout the spinal cord and compare methods to model the hand-grasping task. Additionally, we explore the effect of sample size, data length per participant, and spatial smoothing on activation estimates.

## 2. Materials & Methods

### 2.1 Participants

This study was approved by the Northwestern University Institutional Review Board, and informed consent was obtained for all participants. MRI data were collected from 30 healthy participants (25.9±4.5 years, 11 M). Data were excluded for 4 participants due to excessive image artifacts, an incidental finding, and technical issues with force recordings. All subsequent analyses represent data from the remaining 26 participants (25.8±4.5 years, 9 M).

### 2.2 Data Collection

Spinal MRI data were acquired on a 3T Siemens Prisma MRI system (Siemens Healthcare, Erlangen, Germany) with a 64-channel head/neck coil. A SatPad^TM^ cervical collar (SatPad Clinical Imaging Solutions, West Chester, PA, USA) was positioned around the neck of the participants to increase the homogeneity of the magnetic field around the imaging region. Anatomical and functional scans were acquired before (S1) and after (S2) the administration of a 30-minute acute intermittent hypoxia (AIH) protocol, as part of a larger research study.

#### 2.2.1 AIH Protocol

AIH has been effective at improving motor function in spinal cord injured cohorts^35^. The focus of the broader study was to investigate with MRI the mechanisms of neural plasticity associated with AIH. See Sandhu and Rymer (2021) for a detailed review of AIH in animal models and humans^35^. This AIH protocol was administered after the first MRI session with a HYP 123 oxygen generator (Hypoxico, Inc., New York, NY, USA) in 15 2-minute cycles. Each cycle consisted of brief exposures to a hypoxic gas mixture (9% FiO2, 30-60 seconds), alternating with normal room air (21% FiO2, 60-90 seconds), targeting 85% SpO2 during each bout. The second MRI session occurred 45-60 minutes after AIH. Note, the impact of AIH was not the focus of this work, but it will be discussed in later sections.

#### 2.2.2 Imaging Protocol

A high resolution anatomical T2-weighted scan, covering the brainstem to upper thoracic spine, was acquired with the following parameters: repetition time (TR)/echo time (TE)=1500/135ms, sagittal slice thickness=0.8mm, in-plane resolution 0.39mm^2^, 64 slices, flip-angle (FA)=140°, field-of-view (FOV)=640mm^2^. Spinal cord fMRI scans were collected using a gradient-echo echo-planar imaging (EPI) sequence and ZOOMit selective excitation (TR/TE=2000/30ms, axial in-plane resolution=1mm^2^, axial slice thickness=3mm, 25 ascending interleaved slices, FA=90°, FOV=128×44mm^2^). The ZOOMit acquisition reduces the field-of-view around the spinal cord. The functional acquisition volume was positioned perpendicular to the spinal cord; the bottom of the volume was positioned at the bottom of the C7 vertebral level. Cervical coverage was approximately from the C4 to C7 vertebral level (i.e., approximately C5–C8 spinal cord segments^36^), but the exact coverage varied due to participant height and spinal anatomy. For each scan session, the task paradigm was displayed to participants via a mirror on the head coil, reflecting a monitor placed behind the bore of the magnet. Additional functional scan (1 spinal cord, 2 brain) and a T1-weighted brain anatomical scan were also acquired during the sessions and are not analyzed in this study. The spinal cord T2-weighted and hand-grasp task fMRI scans described here were the first scans acquired during each scan session.

#### 2.2.3 Hand-Grasp Motor Task

A hand-grasping motor task paradigm was designed to elicit cervical spinal cord motor activity. To customize the hand-grasp task paradigm for each participant, MVC was measured prior to each scan session. The participant was seated, and their arm was positioned in 0 degrees of elbow flexion (i.e., fully extended) and the forearm in a neutral position between supination and pronation; they were instructed to grip a Jamar Hand Dynamometer to their maximum force, three times on each side. These values were averaged and input into a custom MATLAB script, which delivered task instructions and real-time force feedback during the scan, targeting 25% MVC (**Fig. 1A**). Two custom, MR-safe, hand-grip devices were designed by affixing two halves of a Delrin rod to an MR-compatible 1-degree-of-freedom load cell (Interface Inc., Scottsdale, AZ, USA). These hand-grips were positioned in the participant’s hands at the start of each scan session. The hand-grasp task paradigm consisted of 24, 15-second left- and right-hand grasps. Right and left grasps were pseudo-randomized such that no more than two consecutive grasps occurred on the same side. Rest periods between each trial were pseudo-randomized to fill the total 10-minute scan duration (300 volumes).

**Figure 1.**
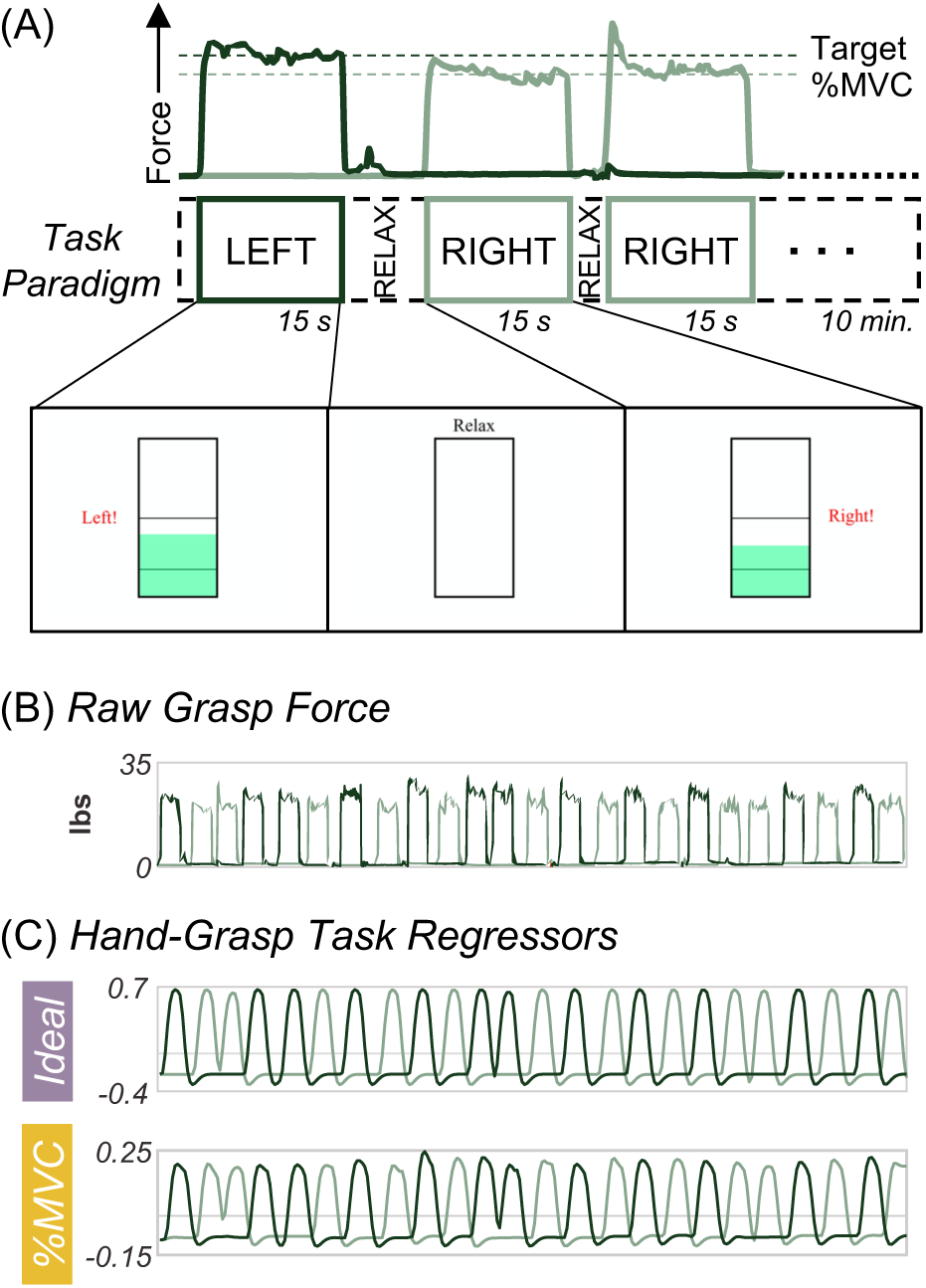
Hand-grasping motor task and task regressors for fMRI modeling. **(A)** First three trials of task paradigm including the grasp force recording to a target %MVC and a representation of the real-time visual feedback (i.e., moving green bar) as the participant is grasping during functional scans. **(B)** The unprocessed grasp force that is used for visual feedback and is recorded. **(C)** The two hand grasp task regressors: Ideal and %MVC. The Ideal regressor was modeled as a binary block design task, convolved with the canonical HRF, and demeaned. The %MVC regressor is the grasp force recording normalized to participant MVC, convolved with the canonical HRF, and demeaned. All panels are from the same example participant.

#### 2.2.4 Physiological Monitorings

Physiological data were collected throughout the fMRI scans including exhaled CO2 and O2 through a nasal cannula, breathing via respiratory belt, and cardiac signal through a pulse transducer on the dorsalis pedis artery of the foot. Pulse could not be recorded from the finger because participants were holding hand-grips. These signals were fed through a Gas Analyzer (CO2, O2 only) and PowerLab and recorded with LabChart (ADInstruments, Sydney, Austrailia). Hand-grasp force data were also recorded through the PowerLab/LabChart system (**Fig. 1B**). The scanner trigger was also recorded through the same system to facilitate alignment of all recordings with fMRI timeseries data.

### 2.3 fMRI Preprocessing Pipeline

All images were first converted from DICOM to NIFTI format (*dcm2niix_afni* ^37^).

#### 2.3.1 Motion correction

2D slicewise motion correction was performed with the Neptune Toolbox^38^ (version 1.211227). Three steps of the toolbox were used to complete motion correction, including the manual definition of a “not cord” mask around the spinal cord and CSF region (#5), computation of a Gaussian weighting mask of the spinal cord, derived from the “not cord” mask (#7), and the application of motion correction (#8). The motion correction algorithm used the Gaussian mask as a weight for each voxel, a temporal median image as the target image for the correction algorithm, and used AFNI^39, 40^ (version 22.0.05) to compute and apply the motion parameters to the data (*3dWarpDrive, 3dAllineate*). Temporal median filtering was de-selected. Motion corrected functional data and slice-wise X and Y motion traces were output.

#### 2.3.2 Registration

A binary spinal cord mask in the functional image space was manually drawn (excluding the top/bottom slices) and is used in several later steps. Segmentation and registration were performed with the Spinal Cord Toolbox^41^ (version 5.3.0). The anatomical T2-weighted image of each subject was segmented (*sct_deepseg_sc*^42^) and then registered to PAM50 template space (*sct_register_to_template*)^43^. The binary spinal cord mask in functional image space and anatomical template warping field were used to inform registration of the motion corrected functional data to the PAM50 template space (*sct_regiser_multimodal*). These warping fields were calculated, but not applied until after subject-level modeling.

#### 2.3.3 Additional Nuisance Regressors

##### Respiratory, cardiac, and CO2

Physiological data were preprocessed in a bespoke MATLAB (MathWorks, Natick, MA, R2019b) script. The results of a peak-finding algorithm were manually verified, and then 8 respiratory and 8 cardiac slicewise RETROICOR regressors were calculated^44, 45^. Only cardiac regressors were calculated for one subject without respiratory belt data. An end-tidal CO2 regressor was also calculated and convolved with the canonical hemodynamic response function (HRF).

##### CSF

A CSF mask was defined by subtraction of the spinal cord mask from the “not cord” mask; high variance (top 20% in each slice) voxels were retained in the mask and the average timeseries within this mask was used to create the slicewise CSF regressor^7, 45^.

##### SpinalCompCor

The “not cord” mask was dilated by 18 voxels. It was then subtracted from the dilation, creating a hollow cylindrical mask, defined as the noise region-of-interest (ROI) outside of the spinal cord/CSF region. Slicewise principal component analysis (PCA) by eigenvalue decomposition was performed in the noise ROI; the first 5 “SpinalCompCor” slicewise PCA regressors were selected^46, 47^.

#### 2.3.4 Modeling the Hand-Grasping Task

The grasping force during scanning was recorded, allowing flexibility in how the motor task is modeled. To compare a standard idealized task model and a task model based on the real-time force recordings, two versions of the hand-grasp task regressors were created for each scan (**Fig. 1C**). For the first regressor, the left and right hand-grasping task were each modeled as an idealized block design and convolved with the canonical HRF (*Ideal*). For the second regressor, the left and right hand-grasping absolute force traces collected during the functional scan were normalized to participant MVC and convolved with the canonical HRF to create task regressors (*%MVC*). Thus, the %MVC regressors capture variation in force targeting across individuals and grasp trials, and the dynamics of each grasping task. The Pearson correlation between the Ideal and %MVC regressors across participants and scans is 0.81±0.03 and 0.81±0.02 for the right and left hand-grasp regressors, respectively.

### 2.4 fMRI Analysis

The following fMRI modeling and analyses were conducted using each the Ideal task regressor and the %MVC task regressor.

#### 2.4.1 Subject-level Analysis

FSL^48^ (version 6.0.3) was used to calculate the subject-level fMRI models (FEAT^49^). The subject-level models contained the left and right task regressors and each of the 25 nuisance regressors described above. FILM (FMRIB’s Improved Linear Model) prewhitening was used; the high-pass filter cutoff was set to 100 seconds and all input regressors were demeaned. Four statistical contrasts were defined: Right Grasp>0, Left Grasp>0, Right Grasp>Left Grasp, and Left Grasp>Right Grasp. Output files were warped from subject functional space to the PAM50 template space. The two “contrast of parameter estimate” (COPE) maps for each of the four statistical contrasts were averaged across the two sessions of each subject (S1 & S2).

#### 2.4.2 Group-level Analysis

A group spinal cord mask was calculated as the consensus region of the co-registered subject-level results, and group-level analyses were performed within this mask, only. For each contrast, a non parametric one-sample t-test using threshold-free cluster enhancement and 5000 permutations was used to calculate group-level activation maps with family-wise error (FWE) rate correction (*randomise*^50^) ^51, 52^. We performed equivalent modeling to look for deactivation, however, no voxels reached the significance threshold.

#### 2.4.3 Assessing the Distribution of Group-level Activation

The localization of activity in the spinal cord was assessed by comparing the distribution of activation in spinal cord ROIs between the ipsilateral and contralateral hemicord, dorsal and ventral cord, and across the rostrocaudal length of the cord. ROIs were defined based on the spinal cord template^43^ and include left and right hemicord masks (defined to exclude a 3-voxel midline between hemicords), dorsal and ventral masks, and spinal cord segments C5-C8. Spinal cord segment masks were defined by thresholding (at 0.02) and binarizing the probabilistic spinal cord segments^53^. The percent of total active spinal cord voxels residing within each ROI was assessed. The distribution of t-statistics was also visualized for each ROI. To additionally quantify the lateralization of activation between the ipsilateral and contralateral hemicords, a laterality index (LI) considering the number of active voxels in each hemicord was defined as LI=(I-C)/(I+C) and ranges from entirely ipsilateral (+1) to entirely contralateral (−1). A ventral-dorsal index was similarly calculated, ranging from -1 (entirely dorsal) to 1 (entirely ventral).

#### 2.4.4 Calculation of Temporal Signal-to-Noise Ratio

Typical temporal SNR (tSNR) maps are derived from resting-state timeseries, which were not acquired in this study. Therefore, tSNR was approximated by for each scan by regressing out any signal variance explained by task or noise confounds. The tSNR maps were warped to the PAM50 template space and averaged across scans and subjects to produce a group average tSNR map. This was performed for both the Ideal and %MVC models. The average tSNR in each spinal cord segment was also calculated.

### 2.5 Analyses of Data Quantity and Smoothing for Activation Mapping

Additional exploration was done into the downstream impact of experimental design and analysis choices on group-level activation results, including sample size, number of fMRI runs (e.g., sessions), and spatial smoothing.

#### 2.5.1 Spatial Smoothing

To apply smoothing, the motion-corrected functional data were smoothed in-plane with a 2×2mm^2^ full width at half-maximum (FWHM) Gaussian kernel. Smoothing was applied using a “restricted smoothing” technique^54^, within a mask of the spinal cord (*3dBlurInMask*). This smoothing kernel was chosen based on recent spinal cord fMRI work^55^. All subject- and group-level analyses were re-calculated with spatial smoothing applied.

#### 2.5.2 Sample Size and Number of fMRI Runs

To test the impact of sample size and data quantity on group-level activation maps, the group-level analyses were repeated for sample sizes of N=14, 18, 22, and 26 subjects for both fMRI runs (S1 & S2). The number of fMRI runs was also tested by repeating the group-level analysis at each sample size with only one fMRI run (S1 only, no averaging of COPE maps). Each of these sample size and fMRI run variations were repeated with and without spatial smoothing and for the Ideal and %MVC models. To summarize, all combinations of the following parameters were computed: task regressor (Ideal, %MVC), smoothing level (no smoothing, in-plane smoothing), number of fMRI runs (S1 only, S1 & S2), and sample size (14, 18, 22, 26).

### 2.6 Impact of AIH

This work is derived from a broader study that was originally designed to test the impact of AIH. Considering these spinal cord fMRI hand-grasp data were collected as a part of this study, a two-tailed paired t-test (S2–S1) was conducted to test whether our spinal cord fMRI methods are sensitive to any potential impact of AIH on motor activation patterns using each of the Ideal and %MVC modeling methods.

## 3. Results

### Motor Activation in the Spinal Cord

Significant group-level spinal cord motor task activation is observed for the hand-grasping task. **Figure 2** presents group-level motor activation mapped for the Ideal and %MVC task regressors for the Right Grasp (R>0) and Left Grasp (L>0) task. Activation appears primarily on the side of the cord ipsilateral to the grasping task. Activation is also present throughout most of the spinal cord segments included in our field of view. Activation in the C8 spinal cord segment is minimal but is increased when using the %MVC regressor as compared to the Ideal regressor. Since the distribution of activation may be impacted by signal quality, the group average tSNR across the spinal cord is shown in **Fig. 3**, which demonstrates the substantial decrease at the C8 segment. The tSNR is systematically lower in the inferior regions of the spinal cord, which may be due to distance from the coil or susceptibility artifacts. Group-level activation is also mapped for Right Grasp>Left Grasp (R>L) and Left Grasp>Right Grasp (L>R) contrasts in **Supplementary Fig. S1**.

**Figure 2.**
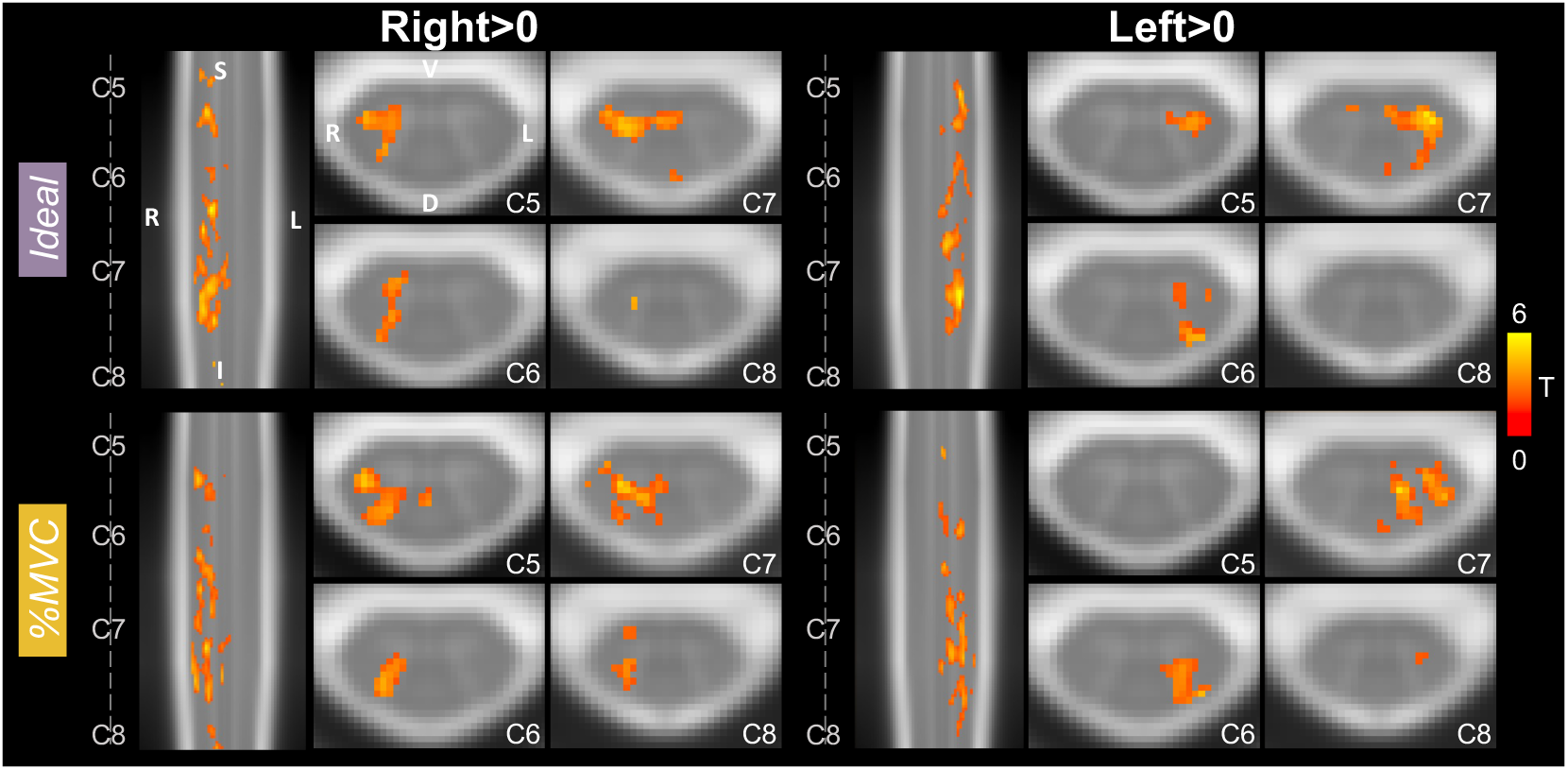
Spinal cord hand-grasp group-level activation maps. Activation maps for each model (Ideal, %MVC) for the Right Grasp>0 and Left Grasp>0 contrasts. Significant t-statistics are shown (p<0.05, FWE-corrected). One representative sagittal slice and 4 axial slices within each spinal cord segment are shown. Probabilistic spinal cord segments are indicated. Note, images are in radiological view.

**Figure 3.**
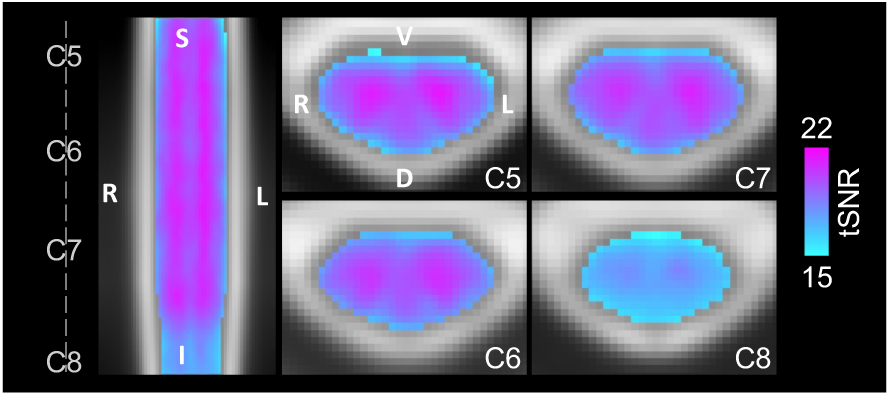
Group average tSNR map after denoising. The tSNR after modeling with the Ideal regressor was calculated for each subject and scan, warped to PAM50 template space, and averaged voxelwise across all subjects and scan sessions. The same representative sagittal and axial slices as Fig. 2 are shown. The average gray matter tSNR across spinal cord segments is C5: 20.14, C6: 20.18, C7: 20.07, and C8: 17.80. The average white matter tSNR across spinal cord segments is C5: 18.30, C6: 18.65, C7: 18.64, and C8: 16.92. See Fig. 4C for a depiction of the spinal cord segment masks tSNR was calculated within. Only one map is shown because there is no visually discernable difference between the tSNR maps after denoising using the Ideal or %MVC model.

### Distribution of Motor Activation

**Figure 4** presents an ROI analysis of active voxels and t-statistics. Activation is primarily constrained to the ipsilateral hemicord for each contrast (**Fig. 4A**). The degree of lateralization is even higher for the R>L and L>R contrasts **(Supplementary Fig. S2)**. The Ideal model laterality indices are 0.977, 0.957, 1.00, and 1.00 for R>0, L>0, R>L, and L>R, respectively. The %MVC model laterality indices are 0.994, 0.996, 1.00, and 1.00 for R>0, L>0, R>L, and L>R, respectively.

**Figure 4.**
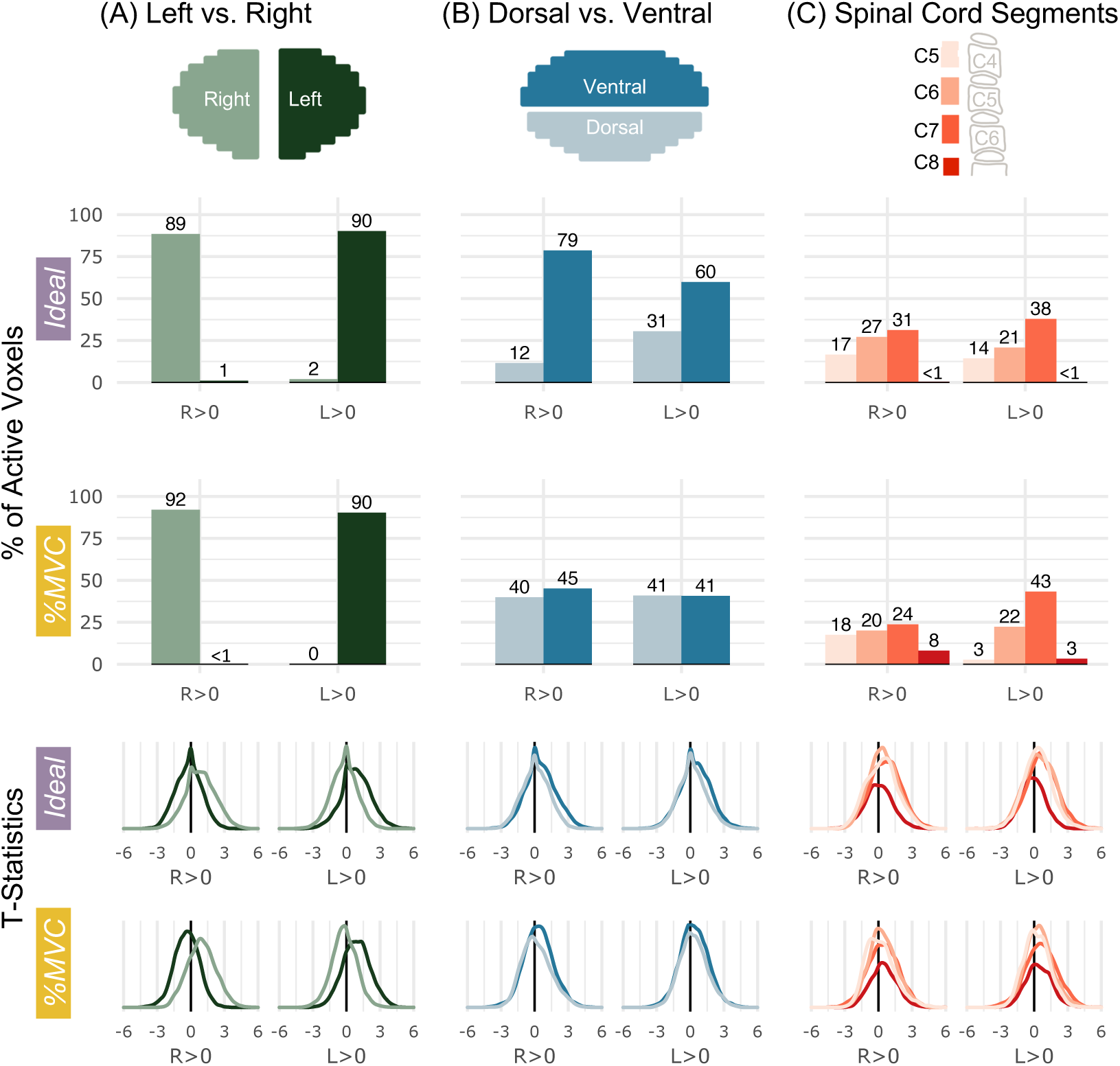
Spatial distribution of significantly active voxels in spinal cord ROIs for Ideal and %MVC models for grasping with the right and left hands. Schematic representations of the ROI masks are shown in the top row. (A) The percent of total active voxels and distribution of t-statistics in the left and right hemicords. The left and right hemicord masks have a 3-voxel midline between the masks. (B) The percent of total active voxels and distribution of t-statistics in the ventral and dorsal hemicords. (C) The percent of total active voxels and distribution of t-statistics that are in spinal cord segments C5-C8. Probabilistic spinal cord segments were thresholded and binarized to create segment masks. Note, active voxels are tallied within these probabilistic masks so there are many voxels not counted as they fall between these segment ROIs. The C8 segment is only partial because it is at the boundary of the field of view. Percentages do not add up to 100 because some active voxels may not fall outside of the ROI mask bounds.

Motor activation is higher in the ventral cord compared to the dorsal cord for the Ideal model, but the distribution is approximately equal for the %MVC model (**Fig. 4B**). The Ideal model ventral-dorsal indices are 0.744, 0.325, 0.230, and 0.087 for R>0, L>0, R>L, and L>R, respectively. The %MVC model ventral-dorsal indices are 0.062, -0.002, 0.202, and 0.189 for R>0, L>0, R>L, and L>R, respectively. The distributions of ipsilateral ventral and dorsal activation along the rostrocaudal axis are also plotted but do not show a discernable preference toward any specific spinal cord segments **(Supplementary Fig. S3)**. The distribution of activation across spinal cord segments is shown in C5, C6, C7, and C8 probabilistic segments (**Fig. 4C**). Overall, activation appears to be the most concentrated in the C7 spinal cord segment. There is an increase in C8 activation in the %MVC compared to the Ideal model for both contrasts.

Activation maps and ROI analysis in discrete spinal cord segments provide an incomplete assessment of the rostrocaudal distribution of activation. **Figure 5** shows the total distribution of active voxels along the length of the spinal cord. For each plot, as expected, there is a peak in the C7 spinal cord segment. There is also substantial activation in other spinal cord segments across the rostrocaudal length of the imaged region. When evaluating the R>L and L>R contrasts, the activation is still most densely, but less distinctly, localized to C7 **(Supplementary Fig. S4)**.

**Figure 5.**
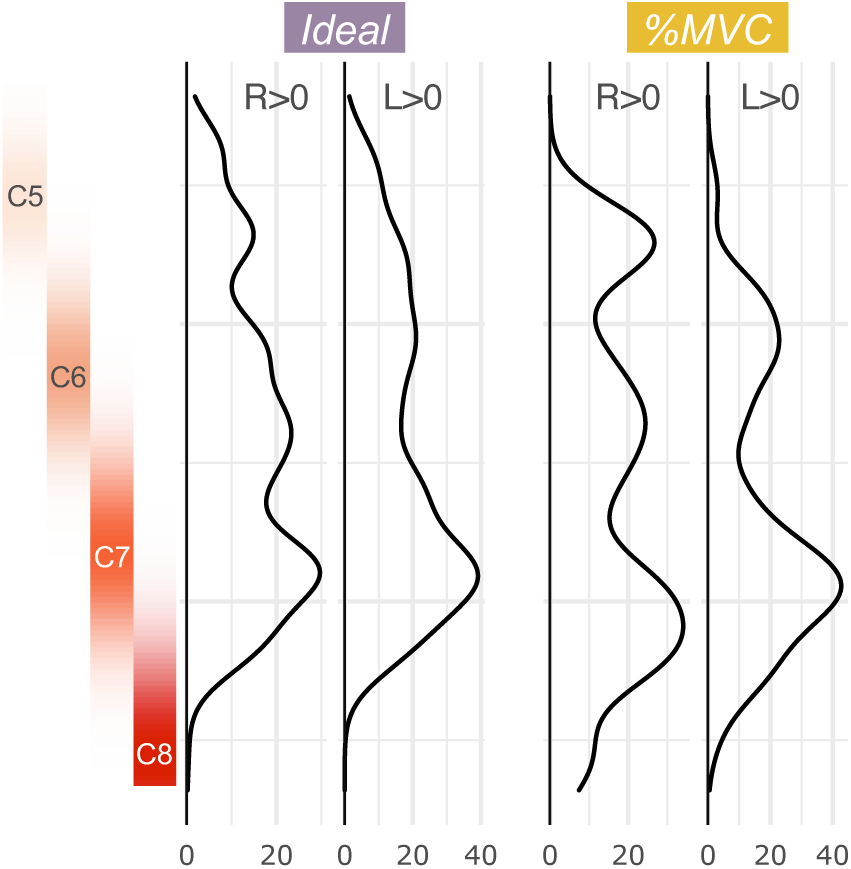
Rostrocaudal distribution of hand-grasp motor activation for the Ideal and %MVC models. The number of active voxels in each axial slice of the group-level activation maps is represented by a density plot for the right and left grasping tasks. Probabilistic spinal cord segments are shown to the left. The activation peak is in the C7 spinal cord segment; however, substantial activation is observed throughout the cord.

### Quantity of fMRI Data and Spatial Smoothing

**Figure 6** shows the effect of sample size, amount of fMRI data, and spatial smoothing on fMRI estimates for the left grasp modeled with the Ideal task regressor. Increasing sample size from 14 to 26 participants noticeably increases the amount and size of activated regions across the rostrocaudal extent of the cord and toward the contralateral hemicord (**Fig. 6A,D**). The addition of in-plane smoothing (2×2mm^2^ FWHM Gaussian kernel) also increases the sensitivity to activation, creating more contiguous clusters of activation at each sample size. Smoothing increases the number of active voxels many times over, especially at lower sample sizes (**Fig. 7**). As anticipated, the group-level parameter estimate (β) peak (summarizing “significantly activated voxels”) tends to decrease as sample size is increased (**Fig. 6B,E**). Doubling the amount of data from one to two 10-minute runs (i.e., S1 only vs. combining S1 & S2) vastly increases the number of active voxels across spinal cord segments, sometimes from zero voxels to hundreds (**Fig. 6C,F**). The number of active voxels for each sample size and smoothing level is also shown in **Supplementary Fig. S5** for S1 only.

**Figure 6.**
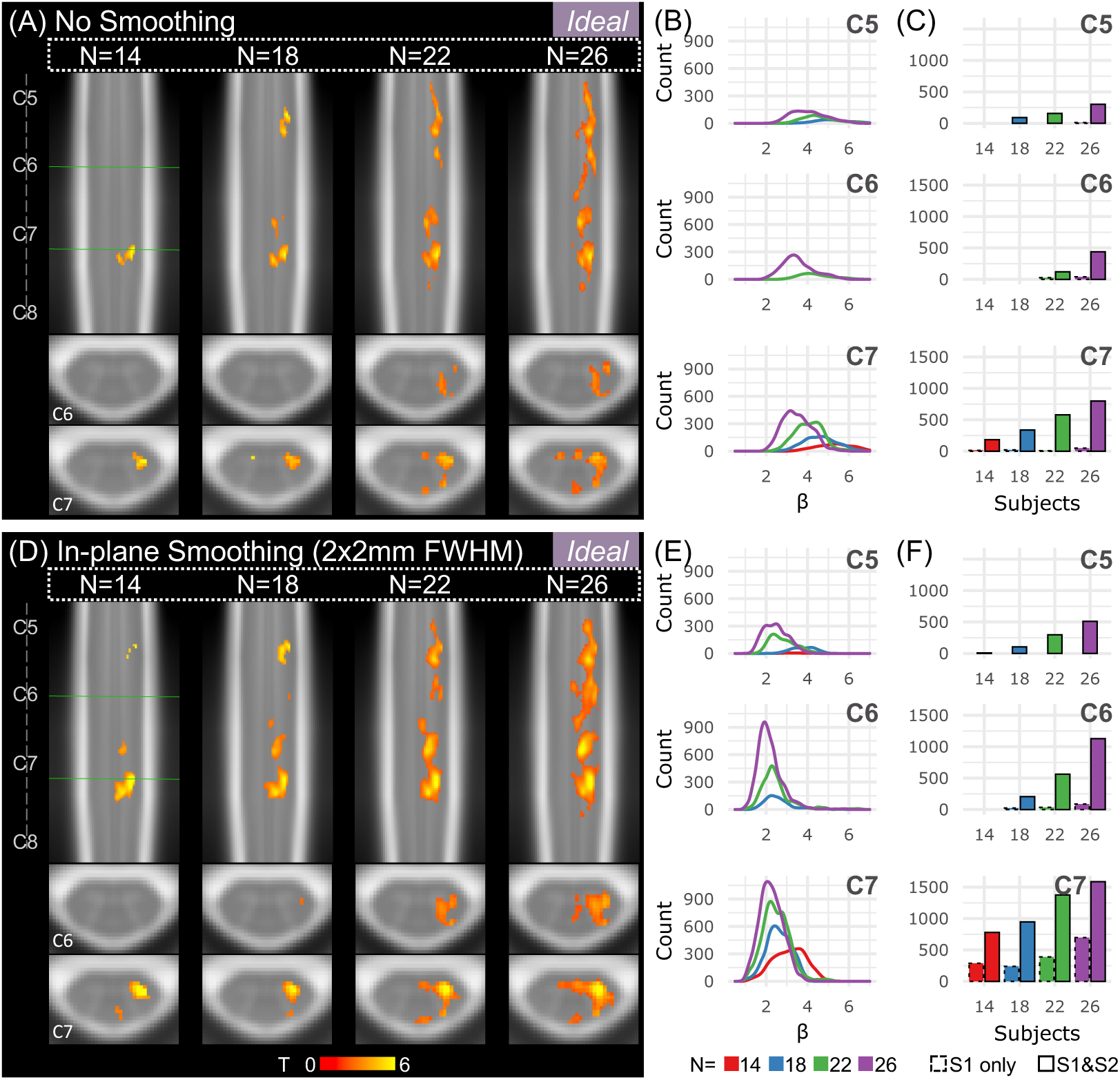
Effects of sample size and spatial smoothing on activation mapping and parameter estimates for the Ideal model, L>0. **(A)** Spinal cord activation map for each sample size: N=14, 18, 22, and 26 participants (S1 & S2). Significant t-statistics are shown (p<0.05, FWE-corrected). **(B)** Density plot distribution of significant parameter estimates for each probabilistic spinal cord segment: C5, C6, and C7 (S1 & S2). **(C)** Number of active voxels in each cord segment for incremental sample sizes using only 1 run (S1 only, 10 min) or 2 runs (S1 & S2, 20 min) for each probabilistic spinal cord segment: C5, C6, and C7. Significant voxels in the C8 segment were very few so are not shown. **(D-F)** Same as A-C for data that were smoothed in-plane within a spinal cord mask using a 2×2mm^2^ FWHM Gaussian smoothing kernel.

**Figure 7.**
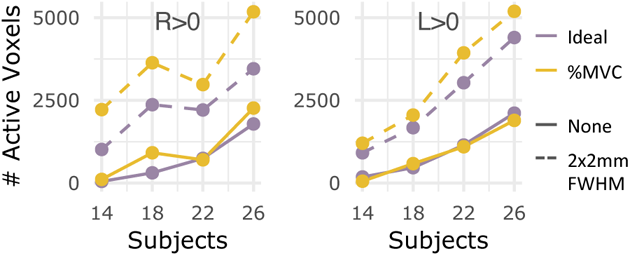
Number of active voxels across sample sizes, with and without smoothing for S1 & S2. The number of significantly active voxels are plotted for sample sizes N=14, 18, 22, and 26 using both fMRI runs (S1 & S2) without smoothing (solid line) and with smoothing (dashed line). Color indicates which task regressor was used for modeling (Ideal, %MVC). The R>0 and L>0 contrasts are shown here.

For each tested regressor (Ideal, %MVC) and statistical contrast (R>0, L>0) combination, Fig. 6 panels B-C and E-F are replicated in **Supplementary Fig. S6**. Notably, for %MVC R>0, activation is already detectable with low sample sizes, so increasing the fMRI data is not as impactful. Additionally, it is sometimes more impactful on the number of active voxels to collect double the amount of fMRI runs per subject (i.e., combining S1&S2) instead of collecting data from nearly double the number of participants (i.e., N=14 vs. N=26). For example, this effect can be seen by comparing the N=14 S1&S2 bar (red, solid outline) to the N=26 S1only bar (purple, dotted outline) in the R>0 C7 spinal cord segment (**Fig. 6C,F**).

### Impact of the AIH intervention

The two scans (S1, S2) were acquired before and after a 30-minute AIH protocol. Comparing scans (S2-S1), there were only 4 significant voxels (p<0.05) observed in the C5 segment for the %MVC model of the L>0 contrast **(Supplementary Fig. S7)**. No other statistical contrasts for either model showed significant voxels. The participant hand-grasp MVC was collected before each scan to calibrate the motor task visual feedback. The average MVC decreased from S1 to S2 by 5.35±9.13 lbs and 4.95±9.51 lbs for the right and left grasps, respectively **(Supplementary Table S2)**; the absence of a widespread difference after AIH may be an effect of fatigue. The minimal impact of AIH on S2 vs. S1 supports our decision to include data from both sessions in our primary analysis.

## 4. Discussion

In this work, we used an isometric left and right unimanual hand-grasping task to map motor task activation with spinal cord fMRI. The experimental design offers flexibility in task modeling between an idealized task regressor (Ideal) and a force-recording task regressor normalized to an individual’s MVC (%MVC). Both methods were able to map robust hand-grasp activation, although there are some inherent differences in the task regressors (i.e., timing, amplitude, trial-to-trial variability), which manifest as differences in the group-level results. Group-level activation from 26 participants was robust and distinctly ipsilateral to the side of the task, and most densely localized to the C7 spinal cord segment.

### Spinal cord hand-grasp activation is highly lateralized

Activation detected from the hand-grasping task is highly lateralized to the ipsilateral hemicord, agreeing with the lateral organization of the spinal cord^56^. Most corticospinal tract fibers, originating in cortical motor areas, cross at the pyramidal decussation, travel down the spinal cord, and synapse in the ventral horn ipsilateral to the side of the movement^57^. Laterality has also been frequently reported in motor task spinal cord fMRI since the methodology’s inception. Our observations agree with those of Weber et al. (2016), also reporting a stronger ipsilateral effect with R>L and L>R contrasts, using a different unilateral isometric task (wrist flexion)^24^.

Laterality of the hand-grasping task is not absolute; some activation is observed in the midline or hemicord contralateral to the task. The intermediate zone is a portion of the spinal cord gray matter between the dorsal and ventral horns, connecting the two hemicords, and surrounding the central canal; it contains spinal interneurons which project within the cord and play a role in reflexes and processing of voluntary motor commands^31, 58^. Motor activity in the contralateral cord has been suggested to be a result of some combined neural processing in the intermediate zone^24^, or commissural interneuron activity^29^. Non-human primate work has suggested the involvement of interneurons in the intermediate zone in grasping, albeit in precision grasping^59^. A similar effect may explain the central and contralateral activation observed in this study.

### The distribution of activation throughout other spinal cord ROIs is less conclusive

Motor activation for the Ideal regressor was more concentrated in the ventral half of the spinal cord, which is expected because motor neuron cell bodies are located in the ventral horns. Each ventral horn is organized into pools of muscle groups, from which signals travel through motor neurons to the peripheral nervous system to innervate the musculature^60^. Hand-grasping primarily involves flexors of the distal musculature. Motor pools for flexors are in the less ventral part of the ventral horn; motor pools for the hand and forearm are in the more lateral part of the ventral horn^61^. However, signal quality and spatial resolution are not high enough to confidently differentiate between specific regions of the ventral horns. In contrast, the distribution of activation for the %MVC regressor is almost equal between the ventral and dorsal ROIs. Dorsal cord activity for both models is likely an effect of sensory or proprioceptive activity associated with performing the task. This supposition agrees with other spinal cord motor task fMRI work which attributes observed dorsal activation to sensory or proprioceptive activity^10, 14, 24, 26, 30^. Higher task effort may lead to increased task-correlated sensory activity if participants find the hand-grips to be uncomfortable at a high force output, which was reported by some participants.

Although white matter activation is not expected, active spinal cord voxels are found in both the spinal cord gray and white matter **(Supplementary Fig. S8).** Spinal cord gray matter is a very small region, and partial volume effects between gray and white matter are inevitable at a 1mm^2^ in-plane resolution. Additionally, to evaluate ROIs at the group-level, fMRI data are registered to template space, a step which commonly uses manual spinal cord segmentations. Variation in manual segmentation does impact registration to template space and group-level activation mapping, although not systematically^62^. Imperfect registration may cause minor misalignment with tissue boundaries. Therefore, we do not feel confident enough in the spatial resolution and registration to distinguish between white and gray matter. Importantly, a gap separates the left-right and dorsal-ventral ROIs, so they are more spatially distinct and trustworthy.

### Activation is most densely localized to the C7 spinal cord segment

Intrinsic and extrinsic muscles that act on the hand are innervated by the median and ulnar nerves of the brachial plexus, originating from the C5-T1 and C8-T1 nerve roots, respectively^63^. In non-human primates, the motor neurons for muscles of the hand and forearm muscles acting on the hand are primarily in C8 and T1^64^. Other spinal cord fMRI studies using tasks similar to hand-grasping (e.g., fist clenching, squeezing a ball) have reported a range of activity peaks throughout the C5-C8 spinal cord segments^9, 10, 15, 19, 25^. Dermatomes C6-C8 would also be particularly relevant for hand-grasping sensory feedback^31^. Therefore, it can be reasonably supposed that hand-grasping activation would be expected throughout multiple spinal cord segments, and to localize to the C7-C8 segments within our field of view.

The peak of activation along the rostrocaudal axis is observed in the C7 spinal cord segment for both the Ideal and %MVC regressor across most statistical contrasts. The group-level consensus mask, accounting for differences in anatomy, covers approximately C5-C8. The signal and tSNR is reduced at more inferior segments due likely to a combination of effects (**Fig. 3**). First, the C8 spinal cord segment is the farthest away from head coil elements in the FOV. Additionally, time-varying respiration-induced differences in the B0 field are also known to impose image artifacts and shifts in spinal cord MRI, maximal at the C7 vertebral level^65, 66^. This overlaps with the C8 spinal level, where we observe the drop in tSNR. Using only the Ideal task regressor, it would be natural to assume that poor signal quality prohibited observation of C8 activity. But, using the %MVC task regressor, activity in this segment is revealed. However, the density of activation at C8 is still minimal. This may still be a result of insufficient signal quality with the coils used, or other image artifacts. Additionally, many active voxels are rostral to the predicted spinal cord segments. This may be due to co-contraction of more proximal upper extremity or postural muscles not directly involved in hand-gripping (e.g., the biceps, which are innervated by C5-C6^31^) or the large inter-individual anatomical variability in the location of spinal cord segments^53^. Future work will include complementary use of electromyography to provide anatomical context for the distribution of spinal cord fMRI activation.

### Sample size and data length affect sensitivity and interpretation of BOLD fMRI activation

Many spinal cord fMRI studies are limited in sample size and/or scan duration, which may limit detection of activation. Weber et al. (2016) investigated the statistical power achieved at incremental time series lengths of 5-30 minutes for their upper extremity motor task, reporting a range of 13-30 subjects required to reach 80% power for a 30-minute time series^24^. The size of our dataset (two 10-minute runs in 26 subjects) enables us to investigate the effects of sample size and data length on regional group-level sensitivity to hand-grasp motor task activation in spinal cord fMRI.

Overall, we demonstrate the improvement in spinal cord fMRI sensitivity associated with increased sample size and number of fMRI runs per subject. Although increasing sample size increases activation observed in the central and contralateral cord, lateralization of motor activation is sustained across sample sizes. Additionally, accounting for two fMRI runs compared to one run considerably increases the amount of detected spinal cord activation. Therefore, it is highly advantageous, and may even be more important, to acquire multiple fMRI runs per subject, as opposed to collecting data from more subjects, if only considering number of active voxels detected.

A sample size of 14 participants would not be uncommon in the spinal cord motor task fMRI literature^3^, in particular, in grasp protocols **(Supplementary Table S1)**. We may only be sensitive to a fraction of true activation with a small sample size. At lower sample sizes, significant activation is concentrated in a smaller region and parameter estimates seem higher, while at higher sample sizes significant activation is observed across the rostrocaudal extent of the spinal cord with lower average parameter estimates. This observation is similar to the effect that Cremers et al. (2017) demonstrate; they simulated a dataset for which a high percentage of brain voxels had weak correlations with a behavioral variable and showed that small samples from this dataset not only underestimated the true spatial extent but also had low power and misleading effect sizes^34^. Our observation of inflated parameter estimates at lower sample sizes is comparable to their finding of misleading effect sizes. This can be dangerous if reports of activation from smaller sample sizes actually seem more compelling (i.e., more focal, larger effect sizes) than results with more diffuse activation patterns. This could be the effect seen here, in which more focal caudal activation agrees better with our a priori hypothesis than the larger spatial extent observed with the full sample size. Increasing the sample size may reveal a true effect of more diffuse activation in the spinal cord, potentially due to task associated sensory activity or co-contraction of muscles extraneous to hand-grasping. We emphasize the need for larger sample sizes and additional fMRI runs for more accurate interpretation of activation distributions in motor task spinal cord fMRI.

### Spatial smoothing increases amount of detected activation

Although the data for the main results of this study were not smoothed, smoothing is common in spinal cord fMRI analyses. Spatial smoothing removes high-frequency spatial noise and boosts SNR, particularly beneficial with small spinal cord fMRI voxels. Conversely, smoothing lowers spatial resolution, especially harmful when hypotheses are specific to gray matter horns.

The unique geometry and anatomy of the spinal cord poses the question of which spatial smoothing method should be used. Standard smoothing with isotropic voxels does not agree with the curved cylindrical shape of the spinal cord. One approach is to straighten the spinal cord, smooth down the centerline, and then un-straighten the spinal cord^41^. Because spinal cord fMRI is commonly acquired with anisotropic voxels, this straightening method has also been adapted to use anisotropic smoothing kernels^45^. On the contrary, smoothing across the rostrocaudal axis may be impacted by the periodicity of anatomical structures along the spinal cord axis; some other studies smooth only in the axial plane^55, 67^. Therefore, we opted for simple 2×2mm^2^ FWHM Gaussian kernel for in-plane smoothing, restricted to within a mask of the spinal cord^54^. This restricted technique is intended to mitigate smoothing in noise from adjacent cardiac-driven CSF pulsations. Unsurprisingly, spatial smoothing increases the amount of active spinal cord voxels and the contiguity of regions of activation, but potentially at the cost of lower spatial precision. If activation is detected without spatial smoothing, this step may not be necessary, especially if the research question concerns small spinal cord tissue ROIs. Spatial smoothing may provide a complementary approach to increase sensitivity alongside increasing sample size and number of fMRI runs.

### Differences in idealized vs. participant-specific models of motor activation

The hand-grasping task has real-time visual feedback, allowing for force targeting that is calibrated to individual MVC. This improves consistency intra-individually across multiple scan sessions and inter individually between subjects. This task design also enables targeting across a range of forces or of graded stimulus levels, as BOLD activation is known to be related to the intensity of a motor task in the brain^68^ and spinal cord^12^. Crucially, recording of the actual force trace during scan acquisition allowed the use of MVC-normalized task regressors. Brain motor-task studies have used recorded force data in fMRI models to account for timing differences by defining task and rest blocks^69^, by creating a force nuisance regressor to remove effects related to the actual achieved force or unintended exertion^70, 71^, or as a force covariate in addition to modeling the task as an idealized regressor^72–74^.

We have identified potential benefits of the controlled task and %MVC task regressor, but how do the Ideal and %MVC models actually differ? The idealized regressor models the task as a standard block design, whereas the %MVC regressor normalizes the absolute force recording to participant MVC. The biggest differences in the input traces are task trial timing and achieved amplitude during grasping. The Ideal task regressor considers the task onset as when the participant is visually cued to begin grasping, whereas the %MVC task regressor has a delay for the participant to begin grasping and reach the target force. The time-to-target delay in task onset between the binary and force trace is 2.62±0.24 seconds and 2.59±0.22 seconds for right and left hand-grasping trials, respectively. Thus, on average, the hand-grasp trials are delayed by more than one TR. Although participants have a consistent force target, there are trial-to-trial amplitude variations around the target %MVC. The achieved %MVC, defined as the average of the middle 10 seconds of a grasp trial, was calculated from the force trace normalized to MVC for each task trial. The achieved %MVC for each grasping trial is 31.42±2.36% and 31.63±2.61%, for the right and left grasping task, respectively. **Supplementary Fig. S9** shows the delay and the achieved %MVC for each participant. These differences in the input traces carry into the regressors. A normalized cross-correlation between the Ideal and %MVC regressors achieves maximum correlation (0.86±0.01 and 0.87±0.01) at 3.69±0.74 seconds and 3.69±0.74 seconds for the right and left hand-grasp regressors, respectively.

After characterizing task regressor differences, there is still uncertainty in whether inter-individual variability in achieved %MVC or intra-individual variability is most responsible for differences in group level activation. To evaluate this question, we created a “Unit %MVC” regressor, which retains intra individual variability in timing and amplitude (i.e., trial-to-trial variability), but is scaled to unit amplitude to eliminate inter-individual variability in task performance and to match the scaling of the Ideal regressor. This regressor should not affect the significance of subject-level activation maps (compared to %MVC), but the statistical contrast of parameter estimates (i.e., COPEs) will be different, which are the group-level model input. The Unit %MVC model activation maps and spatial distribution are very similar to that of the %MVC model **(Supplementary Fig. S10, Supplementary Fig. S11)**. Most notably, the ventral-dorsal distribution is about equal, and the t-statistic density plots are a very similar shape to the %MVC model’s. Additionally, the Dice similarity coefficient (DSC) (*3ddot*) was used to compare the spatial similarity of regions of activation between each of the models (i.e., comparing binary masks of active voxels). The DSC ranges from 0 to 1, where 1 is the most spatially similar. For R>0, the Unit %MVC–%MVC DSC is 0.89 while the Unit %MVC–Ideal DSC is 0.32 (the %MVC–Ideal DSC is 0.34), signifying the high spatial similarity between active voxels in the Unit %MVC and %MVC models. Therefore, it is likely the intra-individual variability is driving the differences between the Ideal model and the %MVC model outputs. The Unit %MVC model may be beneficial in reducing inter individual variability, therefore reducing group-level model bias based on achieved %MVC.

### Limitations

The functional image volume only partially covered the region of the spinal cord expected to be activated in hand-grasping. Had the volume been shifted caudally, the peak of activity would have been expected in the C7-T1 segments. However, signal quality at those lower levels may be too poor, warranting an additional coil to improve signal quality. The decrease in tSNR and its possible attribution to distance from coil elements or respiration-induced susceptibility changes was discussed in a previous section. It is also important to note that the spinal cord is nearby a number of other structures which vary in magnetic susceptibility, such as the vertebrae and intervertebral disks. Spatial distortions to do with gradient-echo EPI sequences are a reasonable and documented concern^4^. Although these effects can be observed in the fMRI volumes, on which we manually segmented the spinal cord during preprocessing, the output of the motion correction and co-registration steps were visually checked and believed to be of sufficient quality. Spatial distribution analyses were performed on ROIs, rather than voxel-specific analyses, which should abate concerns of the downstream impact of spatial distortions on the results of this study. Improvements in field homogeneity and distortion correction strategies would likely improve co-registration and improve the sensitivity of our activation mapping. Slice-specific Z-shimming methods should improve the homogeneity of the magnetic field for spinal cord fMRI^75, 76^ and may be beneficial to employ in future studies. Alternatives to gradient echo imaging methods may further ameliorate distortion effects (e.g., spin-echo methods^77^), but sensitivity of such techniques would need to be evaluated.

Our spinal cord fMRI processing pipeline is very hands on and integrates many software programs. The researcher draws a “not cord” mask and a spinal cord mask for each fMRI dataset. While automation of the “not cord” mask is achievable with currently available tools, segmentation of the spinal cord in functional data is a well-known issue in the spinal cord fMRI community. Although manual spinal cord segmentation may not be a required input to the implemented registration algorithm, we believe it improves the efficacy of co-registration and in turn, the accuracy of group-level results. Additionally, the integration of common functional neuroimaging programs (FSL, AFNI), specialized spinal cord imaging programs (SCT, Neptune), and bespoke analysis scripts can be cumbersome and may decrease approachability of the technique.

The main results of this study are to do with characterizing group-level motor task activation. There is variability in the subject-level activation maps that the group-level analyses average out, leaving the most common signal fluctuation across subjects as group-level activity. This is useful in establishing a normative reference of a motor task, as we aim to do in this study. However, as the field of spinal cord fMRI continues to mature, subject-level analyses of differing pathology presentations, or even healthy spinal cord variability, will become increasingly important.

Lastly, the canonical HRF was used to model the spinal cord BOLD response. It is established that variation in the HRF throughout the brain can have a meaningful impact on analyses^78^. Differences in the temporal dynamics have also been reported in the spinal cord^19^. The use of the canonical HRF, developed for brain fMRI, may lead to imperfect detection of hand-grasping spinal cord motor activity.

## 5. Conclusion

In conclusion, we reported robust detection of hand-grasping activity in spinal cord fMRI. Motor activity was modeled with an idealized task regressor and a regressor normalized to participant maximum grasp force, which differed in timing and amplitude of task trials. They produced activation maps with some spatial variability; lateralization of motor activity was stable. Sample size, number of fMRI runs, and smoothing each improved sensitivity to motor activation, but smoothing lowered spatial precision in activation estimates. Overall, we emphasize the importance of a task that is well-controlled across participants and the impact of collecting additional scans from many participants for spinal cord fMRI studies. Application of individually calibrated tasks in people with motor impairments will be especially beneficial to compare across subjects and between those with and without impairments.

## Acknowledgements

This work was supported by the Center for Translational Imaging at Northwestern University. The authors would like to thank Robert L. Barry for contributions to the development of our data preprocessing pipeline and Sameer Faruquee for contributions to data preprocessing.

