## Supplementary Material for "Spatial distribution of hand-grasp motor task activity in spinal cord functional magnetic resonance imaging"

**Table S1. Analysis methods for previous spinal cord fMRI studies using hand-grasp-like tasks.**

| Authors (Year), Journal | Title | Motor Task | n | Field Strength | Summary of Analysis Methods |
|---|---|---|---|---|---|
| Islam et al. (2019), MRM | Dynamic per slice shimming for simultaneous brain and spinal cord fMRI | bimanual fist clenching at approximately 80% of maximum strength | 9 | 3T | slice-timing correction, RETROICOR, motion correction (3-stage), spatial smoothing (anisotropic), high pass filtering, prewhitening, general linear model, co-register data to standard template, group mixed-effects analysis |
| Giulietti et al. (2008), NeuroImage | Characterization of the functional response in the human spinal cord: Impulse-response function and linearity | unimanual (dominant/ right) rubber bulb squeezing at 1Hz | 7 | 1.5T | motion correction, spatial smoothing, co-register within subject data, functional analysis by a de-convolution analysis and regression analysis (both including baseline and linear trend terms) |
| Ng et al. (2006), NeuroImage | Proton-density-weighted spinal fMRI with sensorimotor stimulation at 0.2 T | bimanual hand-gripping (fist clenching) at 1Hz | 14/ 28 | 0.2T | motion correction, generate statistical maps |
| Stroman et al. (2001), MRI | Characterization of contrast changes in functional MRI of the human spinal cord at 1.5 T | rubber ball squeezing at 1Hz (R/L in separate experiments) | 10 | 1.5T | motion correction, filtering, voxel-wise cross-correlation with task paradigm |
| Stroman and Ryner (2001), MRI | Functional MRI of motor and sensory activation in the human spinal cord | unimanual rubber ball squeezing | 15 | 1.5T | motion correction, filtering, voxel-wise cross-correlation with task paradigm |
| Backes et al. (2001), AJNR | Functional MR imaging of the cervical spinal cord by use of median nerve stimulation and fist clenching | left or right fist clenching at 1Hz | 11 | 1.5T | filtering, voxel-wise analysis of covariates |
| Stroman et al. (1999), MRM | BOLD MRI of the human cervical spinal cord at 3 tesla | unimanual (dominant) squeezing rubber bulb at 1Hz | 25 | 3T | motion correction, filtering, voxel-wise cross-correlation with task paradigm |

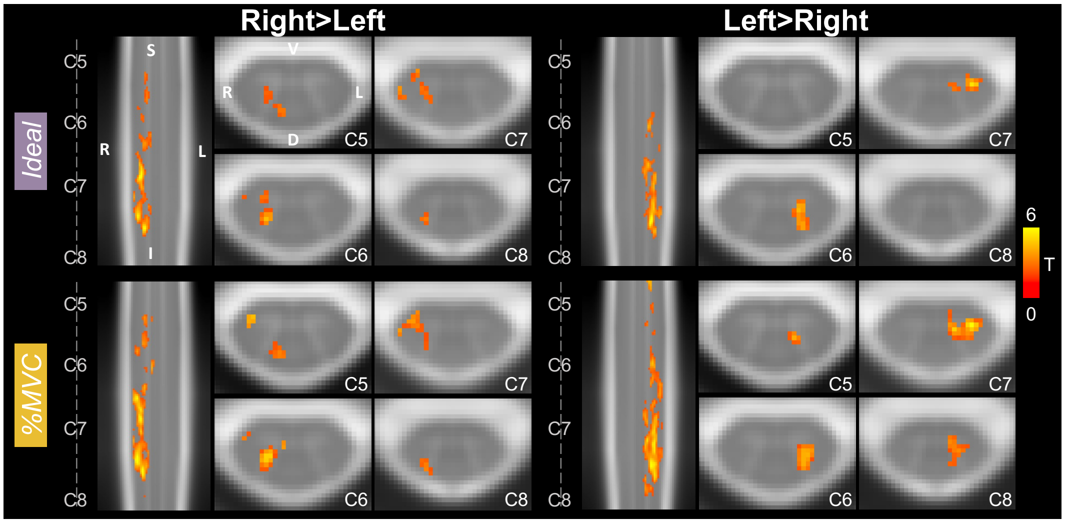


**Figure S1. Spinal cord hand-grasp group-level activation maps.** Activation maps for each model (Ideal, %MVC) for the Right Grasp>Left Grasp (R>L) and Left Grasp>Right Grasp (L>R) contrasts. Significant t-statistics are shown ($p<0.05$, FWE-corrected). One representative sagittal slice and 4 axial slices within each spinal cord segment is shown. Probabilistic spinal cord segments are indicated. Note, images are in radiological view.

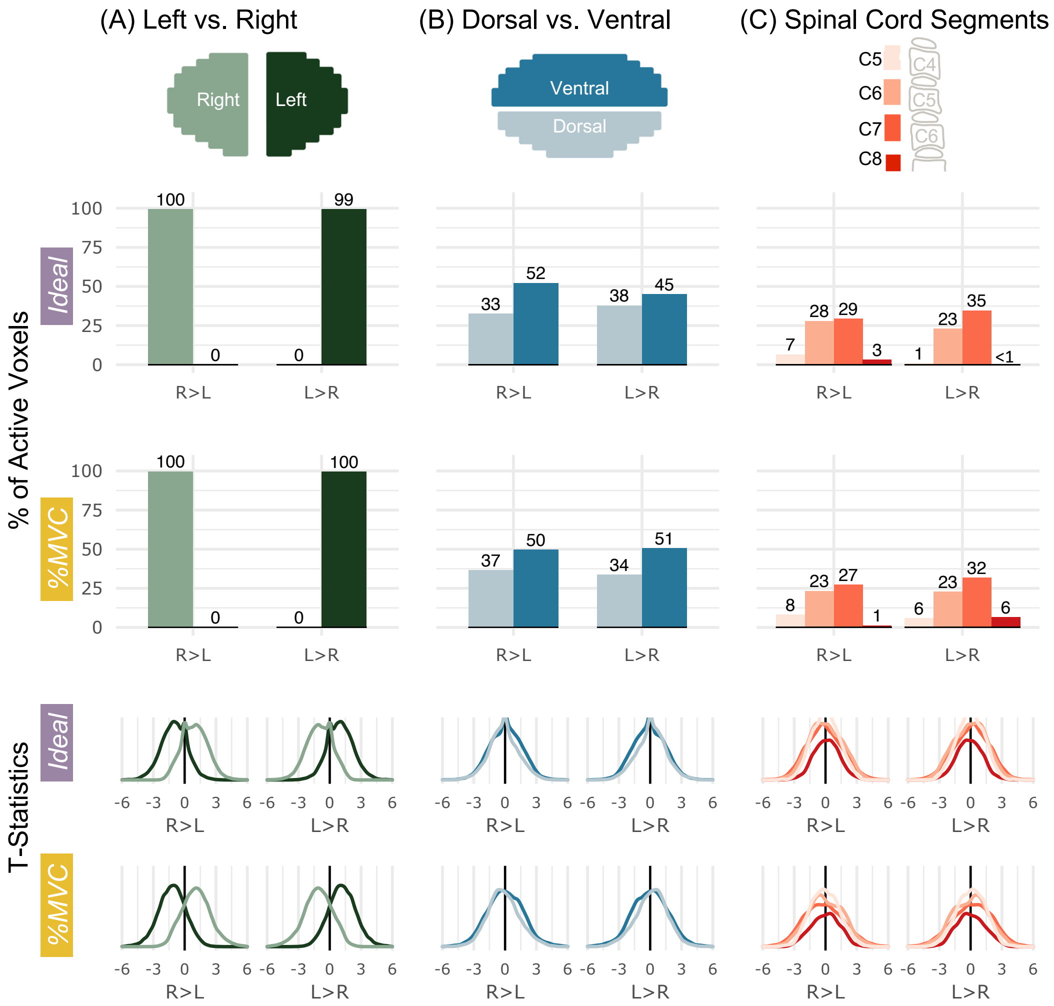


**Figure S2. Spatial distribution of significantly active voxels and t-statistics in spinal cord ROIs for Ideal and %MVC models for R>L and L>R.** Schematic representations of the ROI masks are shown in the top row. **(A)** The percent of total active voxels and distribution of t-statistics in the left and right hemicords. The left and right hemicord masks have a 3-voxel midline between the masks. **(B)** The percent of total active voxels and distribution of t-statistics in the ventral and dorsal hemicords. **(C)** The percent of total active voxels and distribution of t-statistics that are in spinal cord segments C5-C8. Probabilistic spinal cord segments were thresholded and binarized to create segment masks. Percentages do not add up to 100 because some active voxels may not fall outside of the ROI mask bounds. Note, the t-statistics represent a one-tailed t-test; therefore, R>L and L>R distributions are negations of each other.

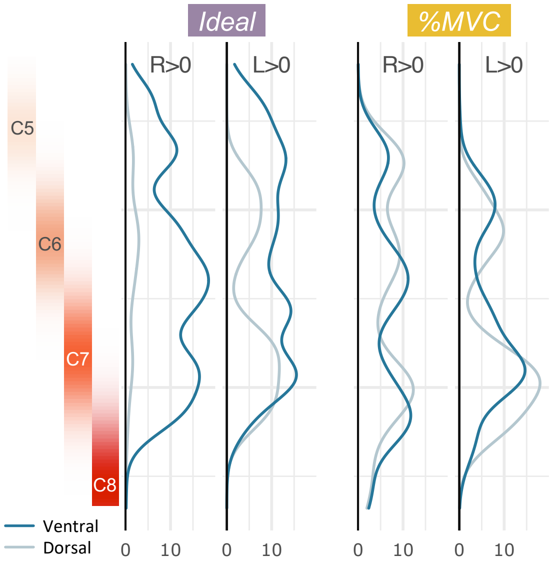


**Figure S3. Rostrocaudal distribution of activation in the ventral and dorsal ipsilateral spinal cord quadrants.** I.e., the right hemicord dorsal and ventral quadrants are shown for the R>0 contrast, and the left hemicord dorsal and ventral quadrants are shown for the L>0 contrast. The number of active voxels in each axial slice are shown. Probabilistic spinal cord segments are shown to the left.

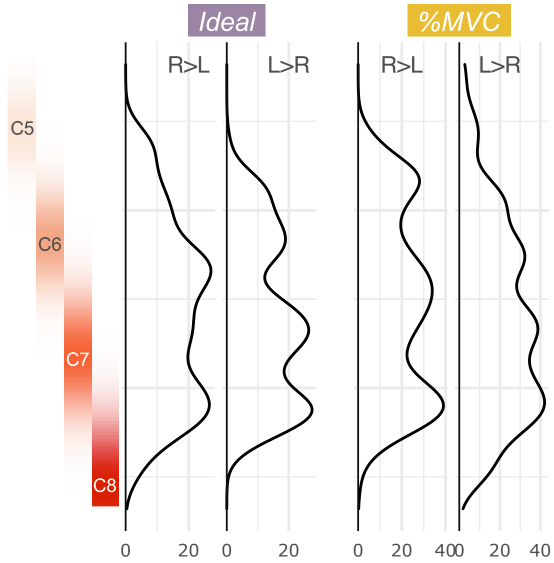


**Figure S4. Rostrocaudal distribution of hand-grasp motor activation for the Ideal and %MVC models for R>L and L>R contrasts.** The number of active voxels in each axial slice of the is represented by a density plot for the right and left grasping tasks. Probabilistic spinal cord segments are shown to the left.

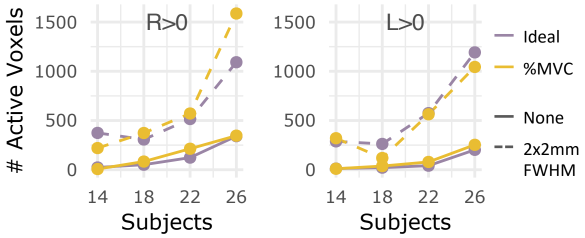


**Figure S5. Number of active voxels across sample sizes, with and without smoothing for S1 only.** The number of significantly active voxels are plotted for sample sizes N=14, 18, 22, and 26 using S1 only without smoothing (solid line) and with smoothing (dashed line). Color indicates which task regressor was used for modeling (Ideal, %MVC). The R>0 and L>0 contrasts are shown here.

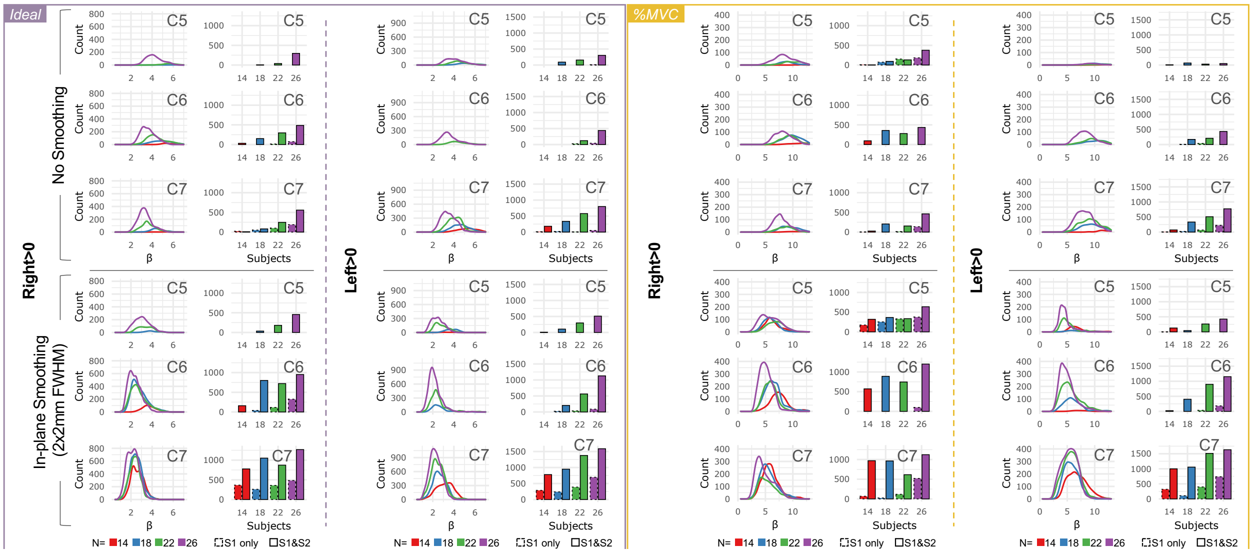


**Figure S6. Effects of sample size and spatial smoothing for Ideal and %MVC models, R>0 and L>0 contrasts, N=14, 18, 22, and 26 sample sizes, and S1 only and S1 & S2.** The Ideal model Left>0 plots are a replicate of **Fig. 5 B-C, E-F** and are shown here for ease of comparison. Shown here across the four sample sizes are density plots of significant parameter estimates and number of active voxels for S1 only and S1 & S2. Note, different y-axis scaling for each contrast.

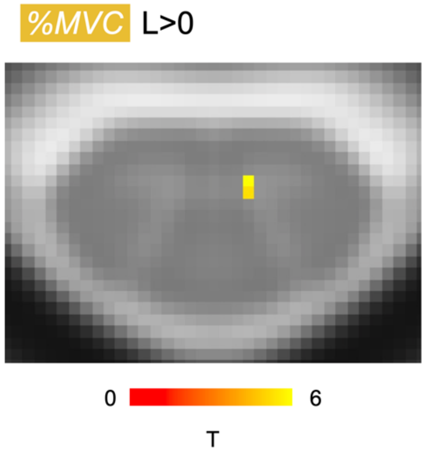


**Figure S7. Effect of AIH on spinal cord hand-grasp activation.** Active voxels represent a significant group-level difference for S2-S1 (two-tailed paired t-test). The only significant voxels observed were for the %MVC L>0 contrast in the C5 spinal cord segment. The Ideal and %MVC models and R>0 and L>0 contrasts were tested.

**Table S2. Left and Right grasp force MVC (lbs) for all subjects and scans.**

| | Scan 1 | | Scan 2 | |
|---|---|---|---|---|
| | **Left** | **Right** | **Left** | **Right** |
| 1 | 123.2 | 120.1 | 120.9 | 118.3 |
| 2 | 64.1 | 65.7 | 68.5 | 66.4 |
| 3 | 60.3 | 53.5 | 54.6 | 53.7 |
| 4 | 82.8 | 96.3 | 78.3 | 97.8 |
| 5 | 73.8 | 70.2 | 72.9 | 70.2 |
| 6 | 112.1 | 132.1 | 99.4 | 121 |
| 7 | 70.2 | 72.2 | 71.8 | 76.6 |
| 8 | 87.5 | 96.5 | 83.3 | 95.2 |
| 9 | 102.9 | 110 | 76.7 | 84.1 |
| 10 | 66.3 | 72.5 | 67.1 | 70.1 |
| 11 | 66.1 | 68.9 | 63.9 | 63.2 |
| 12 | 90.6 | 109.5 | 78.4 | 92.5 |
| 13 | 47.5 | 45.5 | 43.4 | 47.3 |
| 14 | 57.9 | 63.5 | 60.5 | 63.7 |
| 15 | 54.3 | 60.4 | 60.4 | 70.4 |
| 16 | 76.3 | 92.4 | 73.7 | 80.4 |
| 17 | 83.8 | 86 | 70.7 | 79.1 |
| 18 | 78.4 | 76.2 | 86.7 | 71.6 |
| 19 | 62.8 | 64.3 | 44.1 | 54.5 |
| 20 | 54.67 | 65.8 | 49.5 | 64.5 |
| 21 | 39.7 | 41.97 | 36.8 | 36.2 |
| 22 | 43.2 | 56.4 | 36.4 | 45.7 |
| 23 | 84.4 | 100.8 | 83.6 | 98.8 |
| 24 | 61.4 | 65.8 | 50.7 | 53.2 |
| 25 | 76.2 | 76.9 | 46.7 | 47.6 |
| 26 | 76.2 | 77.3 | 78.5 | 90.1 |

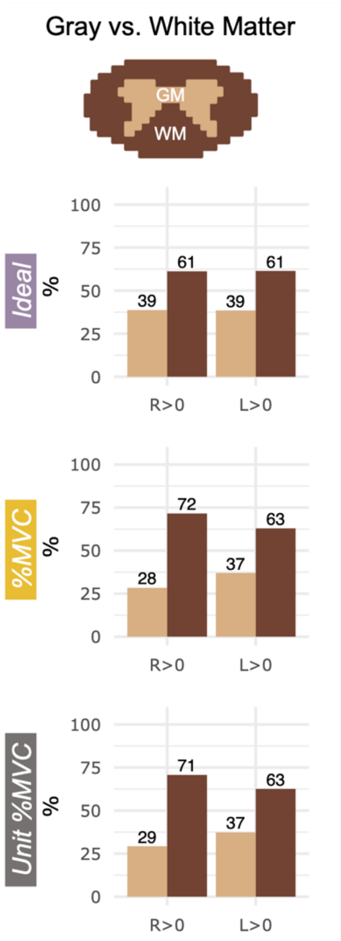


**Figure S8. Distribution of active voxels in white and gray matter ROIs.** The percent of total active voxels for each statistical contrast and the percent that are in each ROI.

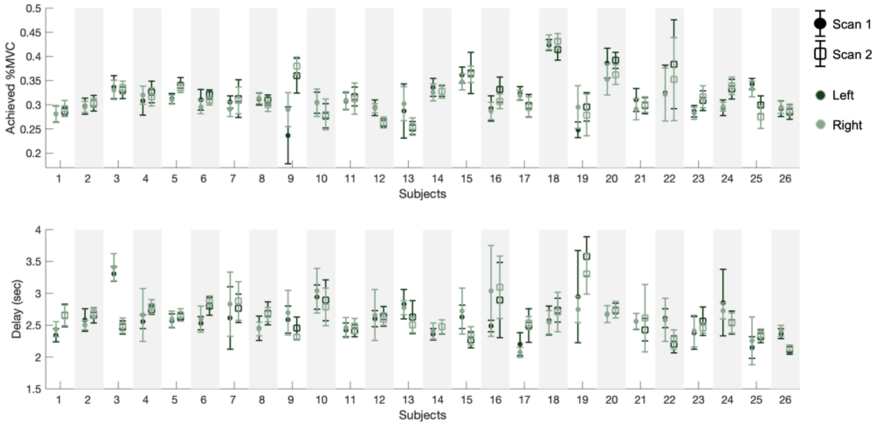


**Figure S9. Variability in achieved %MVC and time to target delay across all subjects and scans.** The achieved %MVC is the force normalized to the participant MVC during the middle 10 seconds of each 15 second trial. The delay signifies the difference in how the Ideal and %MVC task regressors model the task trial timing. Scan 1 (circles) and Scan 2 (squares) are plotted side by side. For each scan, left (dark green) and right (light green) are overlaid.

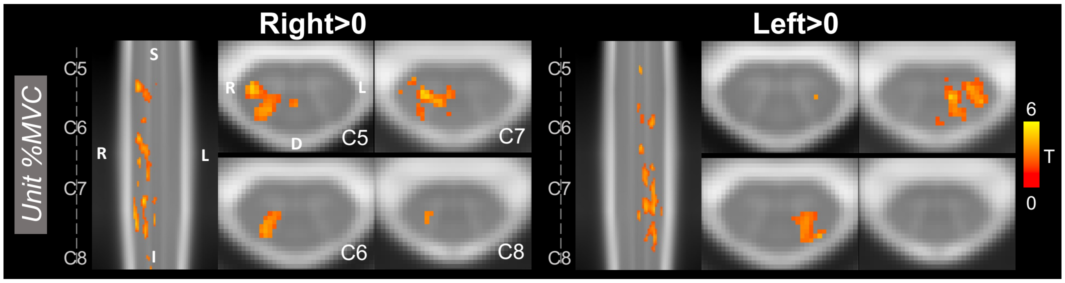


**Figure S10. Unit %MVC activation maps R>0, L>0.** Activation maps for the R>0 and L>0 contrasts. Significant t-statistics are shown ($p<0.05$, FWE-corrected). One representative sagittal slice and 4 axial slices within each spinal cord segment is shown. Probabilistic spinal cord segments are indicated.

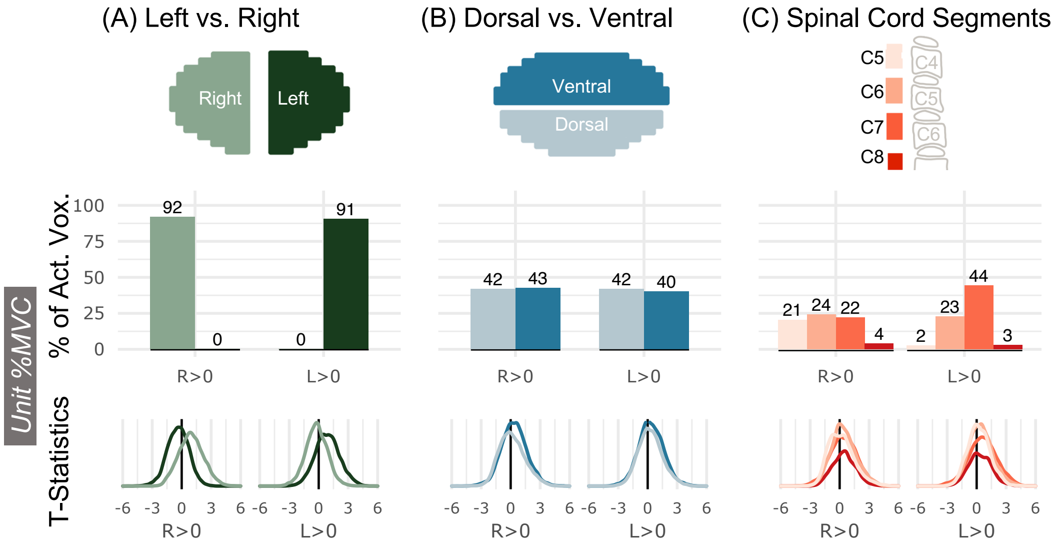


**Figure S11. Spatial distribution of significantly active voxels and t-statistics in in spinal cord ROIs for the Unit %MVC model R>0, L>0.** Schematic representations of the ROI masks are shown in the top row. **(A)** The percent of total active voxels in the left and right hemicords. **(B)** The percent of total active voxels in the ventral and dorsal hemicords. **(C)** The percent of total active voxels that are in spinal cord segments C5-C8. Probabilistic spinal cord segments were thresholded and binarized to create segment masks
